# Trait reward sensitivity and behavioral motivation are associated with connectivity between the default mode network and the striatum during reward anticipation

**DOI:** 10.1101/2025.04.17.649386

**Authors:** James B. Wyngaarden, Akanksha Nambiar, Jeffrey Dennison, Lauren B. Alloy, Dominic S. Fareri, Johanna M. Jarcho, David V. Smith

## Abstract

Individuals vary substantially in their responses to rewarding events and their motivation to pursue rewards. While the ventral striatum (VS) plays a central role in reward anticipation, its functional connectivity with the default mode network (DMN)—critical for self-referential processing and value integration—potentially represents a key mechanism through which trait differences manifest in reward-related behavior. Here, we examine how trait reward sensitivity and state-level behavioral motivation relate to connectivity between the DMN and VS during reward anticipation. Forty-six participants completed the Monetary Incentive Delay task while undergoing fMRI, with trial types reflecting varying levels of reward and loss salience. Behavioral motivation, indexed by reaction time modulation across high-stakes and low-stakes trials, and self-report measures of anticipatory pleasure and reward sensitivity were assessed. Reward sensitivity interacted with anticipatory pleasure to predict behavioral motivation, such that individuals with higher anticipatory pleasure showed weaker behavioral motivation when reward sensitivity was greater, while those with lower anticipatory pleasure showed stonger behavioral motivation when reward sensitivity was greater. Critically, during high-stakes trials, reward sensitivity was associated with stronger DMN-VS connectivity in highly motivated individuals and weaker connectivity in less motivated participants. This moderation effect was consistent across gain and loss contexts, though with distinct directionality patterns. These findings provide novel insights into the neural correlates of individual differences in reward processing, demonstrating that trait reward sensitivity, anticipatory pleasure, and behavioral motivation are associated with distinct patterns of DMN-VS interactions during reward anticipation. These findings highlight the importance of considering motivational context when investigating reward-related neural mechanisms.

## 1. Introduction

Reward processing fundamentally relies on coordinated activity between distributed brain networks that integrate motivational, evaluative, and self-referential information to guide goal-directed behavior (Haber & Knutson, 2010; Pessoa, 2015; Smith & Delgado, 2015). According to Gray’s Reinforcement Sensitivity Theory, individuals exhibit stable differences in their responsiveness to rewarding stimuli, with these trait-level differences reflecting underlying variations in brain systems that process incentive information (Gray, 1970; Gray & McNaughton, 2000). However, contemporary network neuroscience perspectives suggest that reward processing emerges from dynamic interactions between multiple neural systems rather than isolated regional activation (Bassett & Sporns, 2017). Specifically, effective reward processing requires integration between regions that evaluate incentive salience (such as the ventral striatum) and networks supporting self-referential processing (such as the default mode network [DMN]), associated with contextualizing rewards within personal goals and experiences (Andrews-Hanna et al., 2014; Smith et al., 2025). Recent evidence demonstrates that core DMN regions overlap extensively with the brain’s valuation system and serve as integrative hubs for subjective value computation and goal-directed behavior (Smith et al., 2025). This network integration perspective suggests that individual differences in reward sensitivity may be best understood as differences in how reward-related networks coordinate their activity during incentive processing.

Although trait reward sensitivity represents a relatively stable individual difference, its translation into goal-directed behavior depends on state-level motivational processes that respond dynamically to environmental incentives (Brehm & Self, 1989). Motivational intensity theory posits that effort allocation reflects both underlying sensitivity to incentives and perceived task demands, with behavioral motivation—indexed through reaction time modulation—capturing the mobilization of cognitive and motor resources proportional to incentive salience (Locke & Braver, 2008). This approach is supported by evidence that reaction time differences to reward targets reflect individual variation in relative motivation, with nucleus accumbens activation mediating these motivational differences even in the absence of explicit choice (Clithero et al., 2011). This relationship is further complicated by individual differences in anticipatory pleasure capacity, where anhedonia is associated with reduced motivation and constitutes a core feature of depression (Treadway & Zald, 2011). Importantly, anticipatory pleasure capacity may fundamentally alter how trait reward sensitivity translates into motivated behavior, with individuals showing different trait-behavior coupling patterns depending on their anticipatory systems (Pizzagalli et al., 2005; Cho et al., 2016). This suggests that anticipatory pleasure serves as a moderating factor in the dynamic relationship between stable reward sensitivity traits and context-dependent motivational states.

At the neural level, the ventral striatum (VS) serves as a central hub integrating motivational signals from diverse cortical and subcortical regions during reward processing (Sescousse et al., 2013). Emerging evidence suggests that functional connectivity between the VS and other brain networks may be equally important as regional activation for understanding individual differences in reward processing (Camara et al., 2009). The default mode network (DMN), encompassing regions involved in self-referential thinking, autobiographical memory, and personal goal representation, has been particularly implicated in reward-related integrative processes (Buckner & Carroll, 2007; Acikalin et al., 2017; Smith et al., 2025). From a network neuroscience perspective, VS-DMN connectivity during reward anticipation may represent a critical mechanism through which individuals integrate external incentive information with internal representations of self-relevant goals and values, thereby translating raw incentive salience into personally meaningful motivational responses (Utevsky et al., 2014). This connectivity pattern should vary systematically with individual differences in both trait reward sensitivity and state-level motivational factors. In this study, we operationalize behavioral motivation as a task-specific, state-dependent measure indexed by reaction time modulation in response to trial incentive salience, distinguishing it from trait-level motivational tendencies.

Despite growing recognition of network-level interactions in reward processing, previous research examining VS-DMN connectivity has yielded inconsistent findings, with some studies reporting enhanced connectivity in reward-sensitive individuals whereas others suggest reduced connectivity under high motivational demands (Posner et al., 2016; Saris et al., 2020). These discrepancies may arise from failure to account for individual differences in both trait-level reward sensitivity and state-level motivational factors that could systematically modulate VS-DMN interactions. If VS-DMN connectivity reflects integration of external incentive information with internal self-referential processes, then connectivity strength and direction should depend on both underlying reward sensitivity and current motivational state in response to specific incentive contexts (Knutson & Greer, 2008). Furthermore, different incentive types—gains versus losses—may recruit distinct integration patterns, as approach and avoidance systems engage partially overlapping but functionally distinct neural circuits (Elliot, 2006). A comprehensive understanding of VS-DMN connectivity requires examining how stable individual differences interact with dynamic motivational states across different incentive contexts.

To address these gaps, the current study investigates how trait reward sensitivity, state-dependent behavioral motivation, and anticipatory pleasure capacity jointly influence VS-DMN connectivity during reward anticipation. Using an fMRI-based Monetary Incentive Delay task, we examine how individual differences in reward sensitivity interact with task-driven motivational responses across varying reward and loss salience. We first examine how anticipatory pleasure capacity moderates the translation of trait reward sensitivity into behavioral motivation, testing whether individual differences in anticipatory pleasure shape the RS-behavior relationship. Second, we assess whether reward sensitivity predicts distinct patterns of VS activation during reward anticipation, establishing basic striatal response patterns to incentive salience. Third, we test our primary hypothesis that the relationship between reward sensitivity and VS-DMN connectivity will be systematically modulated by behavioral motivation, such that highly motivated individuals will show different connectivity patterns compared to less motivated individuals. Critically, we propose that DMN-VS connectivity represents the key neural integration mechanism through which reward sensitivity and motivational state interact to guide reward-related behavior. By integrating behavioral and neural perspectives within a unified theoretical framework, this study aims to resolve previous inconsistencies while advancing understanding of how individual differences shape network-level reward processing mechanisms.

## 2. Materials and Methods

### 2.1 Participants

This dataset is available as OpenNeuro Dataset 4920 (Smith et al., 2024), and it is composed of neuroimaging data from 59 participants who completed four tasks involving social and nonsocial reward processing. The pre-registration (https://aspredicted.org/PQA_WPB) describes the goal to collect data from 100 participants (18-22), wherein we acquired data from 60 participants due to constraints imposed by the COVID-19 pandemic. As per pre-registered criteria, fourteen of the 60 participants who completed the study were excluded from analyses due to their failure to respond during behavioral tasks (>20% missing responses; N=4), incomplete data (N=4; failure to complete survey data or missing behavioral data due to technical issues), or poor image quality (N=6). Image quality was defined using the fd_mean and tSNR values from MRIQC. Participants were excluded for fd_mean values greater than 1.5 times the interquartile range, per the distribution from neuroimaging data of otherwise eligible participants. This resulted in the final set of 48 participants (mean age: 20.45 yrs, SD: 1.89 yrs; 22.7% male of which 57% white, 34% Asian, 9% other-2 Black/African American, 1 black and white, 1 Indian). For a description of deviations from the pre-registration, see Supplementary Methods 1.

Participants were recruited via the Temple University Psychology and Neuroscience Department participant pool, and from the surrounding community via flyers and online advertisements. Participants were paid $25 per hour for fMRI and $15 per hour for behavioral tasks, and received bonuses based on their decisions on other neuroeconomic tasks (not reported here), resulting in a total payment of $140 to $155. In addition, participants recruited from the university pool also received research credit for their participation.

### 2.2 Procedure

All methods were approved by the Temple University IRB. Prospective participants were identified based on their responses to an online screener questionnaire, which assessed RS using the Behavioral Activation Subscale (BAS; Carver & White, 1994) and the Sensitivity to Reward subscale (SR; Torrubia et al., 2001). A sum was calculated for each subscale. Sums were assigned to a quintile that reflected low to high levels of RS across the distribution of possible scores. We used methods consistent with our prior work (e.g., Alloy et al., 2012) to ensure participants were responding truthfully and attentively. Only participants with scores within +/-1 quintile on both subscales were eligible for the study (no exclusions were made based on this criteria). At the in-person visit, we confirmed that eligible participants were free of major psychiatric or neurologic illness and MRI contraindications. Prior to MRI scanning, all participants underwent safety screening including verification of MRI compatibility to ensure data quality and participant safety.

In order to probe behavioral motivation, participants were subjected to the Monetary Incentive Delay Task (MID) (Knutson et al., 2000). During the task, participants respond to a stimulus in order to either gain money or avoid losing money (Fig. 1). There are 5 conditions, corresponding with the 5 different shapes in the top panel (Large Loss, Small Loss, Neutral, Small Gain, Large Gain). Precisely, the value of money presented on the screen was (-$5, -$1, $0, +$1, +$5). During the cue phase, participants see a shape which indicates the money at stake for the current trial. They are then presented with an ISI (inter-stimulus interval) period until a white target square appears. Once the square appears, they have 1 second to respond. If they respond in time, they win the gain trials, acquiring money, and in loss trials, they avoid losing money. In contrast, if they do not respond quickly, they lose the trial; i.e., in gain trials they don’t win money, and in loss trials they lose money.

**Figure 1.**
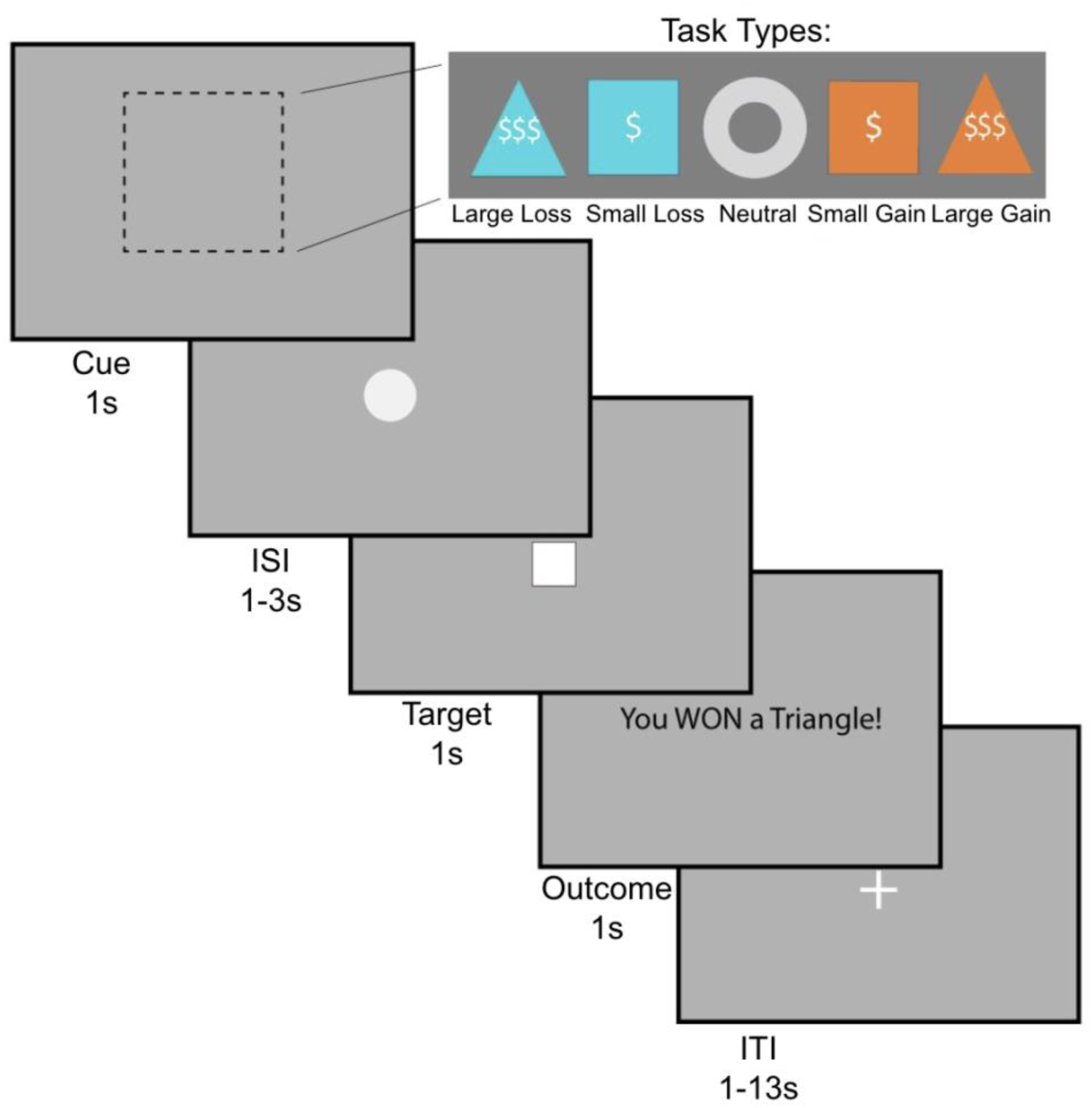
The monetary incentive delay (MID) task. During the cue phase, participants see a cue indicating the trial type: large gain, small gain, large loss, small loss, or neutral (1s). Then, after a jittered inter-stimulus interval (1-3s), the target square appears (1s), during which time participants must respond with a button press. This is followed by an outcome phase (1s). Trials are separated by a jittered interval (1-13s). Figure adapted from Smith et al., 2024 with permission.

The anticipation regressor was defined as the interval from cue onset to target onset, modeling neural activity related to the expectancy of potential monetary outcomes prior to the motor response. This period directly precedes the RT measurement window, making it a key neural correlate of the motivational processes that influence speed of response during the task. Thus, the MID task analyzes people’s behavior based on reaction time or how quickly people respond when the cue appears.

### 2.3 Individual difference measures

Reward sensitivity (RS) was assessed using a composite measure derived from the z-scores of two self-report scales: the Behavioral Activation System (BAS) scale (Carver & White, 1994) and the reward subscale of the Sensitivity to Punishment and Sensitivity to Reward Questionnaire (SPSRQ; Torrubia et al., 2001). The BAS scale is a 20-item questionnaire designed to measure individual differences in sensitivity to reward and approach motivation. The SPSRQ reward subscale consists of 24 items assessing sensitivity to reward in specific contexts. Both the BAS and SPSRQ reward subscale have been established as reliable and valid measures of trait reward sensitivity (Alloy et al., 2006; Alloy et al., 2012). Additionally, we included the Temporal Experience of Pleasure Scale (TEPS; Gard et al., 2006), an 18-item measure designed to assess distinct components of pleasure experience. We employed the TEPS anticipatory subscale because it specifically targets anticipatory pleasure processes that align with the MID task’s focus on reward anticipation rather than consumption. TEPS has established psychometric properties and clinical validity for capturing individual differences in anticipatory pleasure capacity relevant to reward processing. Both the anticipatory (TEPSa) and consummatory (TEPSc) subscales were examined. The two subscales showed a moderate positive correlation (r = .55, p < .001), indicating they measure distinct but related components of pleasure experience. In behavioral analyses, consummatory pleasure was included as a covariate to assess the unique contribution of anticipatory pleasure to reward processing during anticipation-focused tasks.

We also aimed to examine individual differences in behavioral motivation during the MID task. We operationalized behavioral motivation as the extent to which reaction times (RTs) were influenced by reward salience within the task context, indexing it through three RT contrasts (Fig. 2A). This task-specific operationalization captures context-dependent motivational responses rather than stable dispositional motivational tendencies. First, HS>LS (high stakes vs. low stakes) captures the overall pattern of RT differentiation across trial types, such that individuals with greater behavioral motivation responded more quickly to high-stakes trials (large gain: +$5; large loss: -$5) compared to low-stakes trials (neutral: $0; small gain: +$1; small loss: -$1). To quantify this, we fitted a second-degree polynomial to each participant’s RTs across all five trial types, extracting the quadratic coefficient as HS>LS. A more negative HS>LS value indicates a stronger pattern of faster RTs for high-stakes trials and slower RTs for low-stakes trials, reflecting greater behavioral motivation, whereas a less negative, near-zero, or positive value indicates weaker RT differentiation or faster RTs for low-stakes trials, reflecting lower motivation. Negative values are expected for individuals with strong behavioral motivation, though positive or near-zero values may occur due to individual variability in task engagement, arousal, or response strategies. Second, we examined specific contrasts: LG>N (large gain vs. neutral) captures motivational responses to large rewards, and LL>N (large loss vs. neutral) captures motivation to avoid significant losses. Similarly, for LG>N and LL>N, more negative values indicate faster RTs for large gain or loss trials relative to neutral trials, reflecting higher motivation, whereas less negative, near-zero, or positive values indicate lower motivation, consistent with individual differences in response patterns. These complementary approaches allowed us to assess both the overall pattern of behavioral motivation (via HS>LS) and its sensitivity to specific reward or loss conditions (via LG>N and LL>N). Notably, avoiding a large loss is itself a highly motivating outcome, as individuals often treat potential losses as psychologically equivalent to missing out on a comparable gain (Kahneman & Tversky, 1979). This aligns with evidence that loss aversion engages reward-processing circuitry, reinforcing the idea that both large gains and large losses can drive motivated behavior.

**Figure 2.**
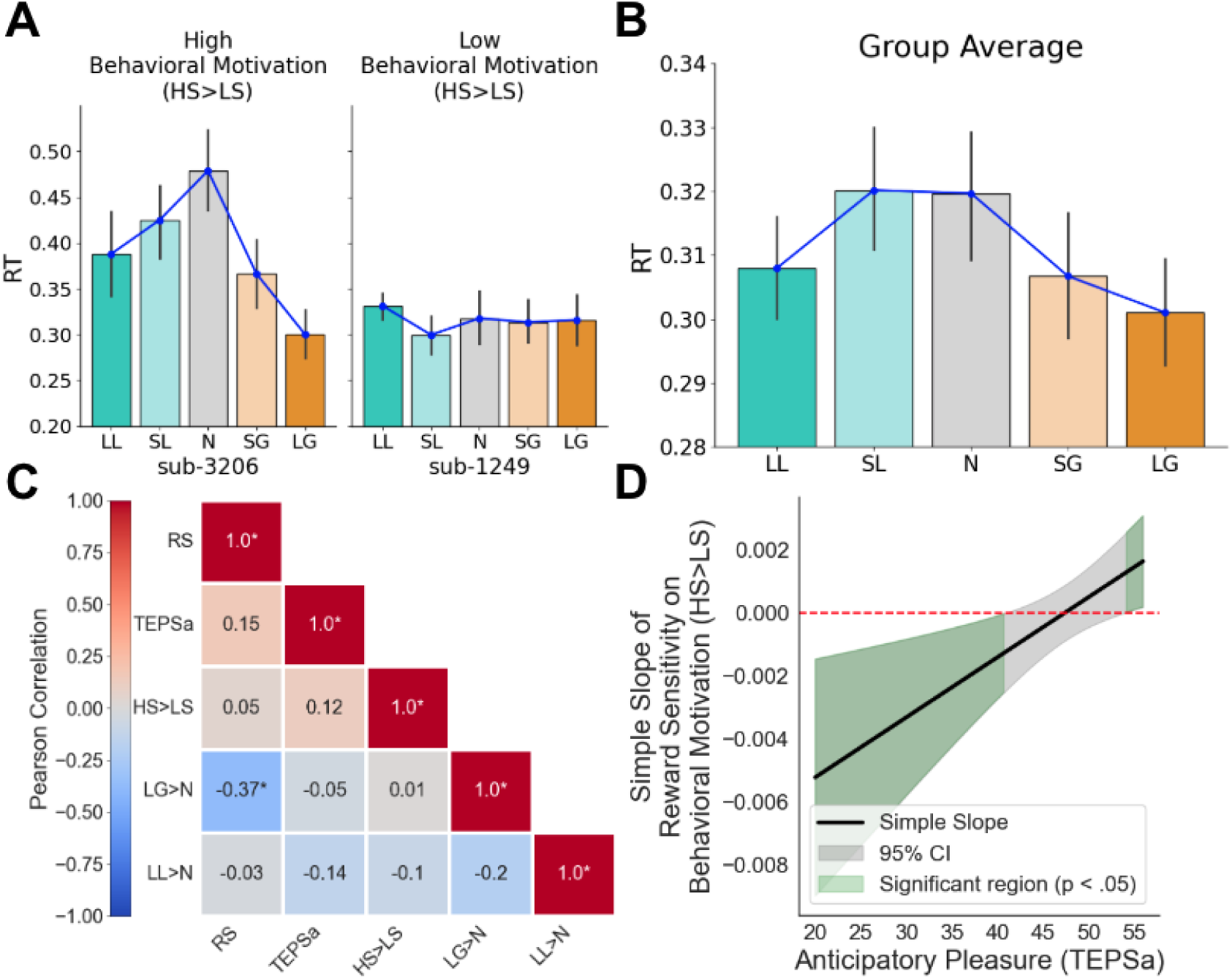
Anticipatory pleasure (TEPSa) modulates the relationship between Reward Sensitivity (RS) and Behavioral Motivation (HS>LS). (A) Examples of high (characterized by faster RTs for high stakes trials relative to low stakes trials) vs. low (no difference in RT between trials) behavioral motivation. (B) Behavioral motivation across the sample. On average, participants responded faster for large gain and loss trials than for neutral trials, demonstrating behavioral motivation on reward-salient trials. (C) Correlation heatmap of variables of interest, where LG>N refers to the contrast in RTs for Large Gains vs. Neutral and LL>N refers to the contrast in RTs for Large Losses vs. Neutral. (D) Johnson-Neyman plot showing the simple slope of reward sensitivity on behavioral motivation (HS>LS) across anticipatory pleasure (TEPSa). Green regions indicate significant slopes (p<.05) at low TEPSa (low anticipatory pleasure) and high TEPSa (high anticipatory pleasure); the gray region indicates non-significant slopes.

### 2.4 Neuroimaging Data Acquisition and Preprocessing

These data were collected as part of an overarching project that has been described in a previous publication (for full details see Smith et al., 2024). Images were acquired using a simultaneous multi-slice (multi-band factor = 2) gradient echo-planar imaging (EPI) sequence (240 mm in FOV, TR = 1,750 ms, TE = 29 ms, voxel size of 3.0 x 3.0 x 3.0 mm^3^, flip angle = 74, interleaved slice acquisition, with 52 axial slices). Neuroimaging data were converted to the Brain Imaging Data Structure (BIDS) using HeuDiConv (Halchenko et al., 2024). Results included in this manuscript come from pre-processing performed using fMRIPrep 20.2.3 (Esteban et al., 2019), which is based on Nipype 1.4.2 (Gorgolewski et al., 2011, 2016). Following fMRIprep, we used FSL to apply a 5 mm smoothing kernel. Additional MRI acquisition and preprocessing parameters are described in the supplementary materials.

### 2.5 FMRI Analyses

Neuroimaging analyses used FSL version 6.0.0 (Smith et al., 2004; Jenkinson et al., 2012) with individual-level general linear models and local autocorrelation correction (Woolrich et al., 2001). The activation model focused on brain responses during the anticipation phase using seven task-based regressors: anticipation of large gains, small gains, large losses, small losses, neutral trials, hits, and misses (jittered 2000-4000 ms). This anticipation-focused approach isolated neural activity associated with reward anticipation rather than outcome. All models included six motion parameters (rotations and translations), the first six aCompCor components, and framewise displacement as covariates of no interest, with high-pass filtering (128s cut-off) applied using discrete cosine basis functions.

The connectivity model examined task-dependent DMN-VS interactions using network psychophysiological interaction (nPPI) analysis (Friston et al., 1997; O’Reilly et al., 2012). The first seven regressors were identical to the activation model. The DMN and nine additional networks were defined based on prior work (Smith et al., 2009). Network time courses were extracted with a spatial regression component of the dual regression approach (Filippini et al., 2009; Nickerson et al., 2017) and entered into a model with the seven task regressors from the activation model described above. PPI regressors were formed by multiplying each of the seven task regressors by the DMN regressor, yielding a total of 24 regressors. This approach examined incentive-salience-dependent changes in DMN connectivity with bilateral VS (Oxford-GSK-Imanova atlas; Tziortzi et al., 2011).

For group-level analyses, we conducted two sets of pre-registered ROI-based tests. First, VS activation during incentive anticipation was examined by extracting BOLD signal from bilateral VS and conducting linear regressions for incentive salience contrasts (HS>LS, LG>N, LL>N) with reward sensitivity (RS), anticipatory pleasure (TEPSa), and their interaction as predictors. Second, DMN-VS connectivity was examined using nPPI-derived connectivity estimates regressed onto RS, behavioral motivation (matching RT contrasts to connectivity contrasts), and their interactions, as well as models including TEPSa interactions.

Multiple comparison corrections were applied using Tukey’s HSD for pairwise comparisons and Bonferroni correction for family-wise error control across incentive salience contrasts (detailed procedures in Supplementary Methods). All statistical analyses were performed using R (version 4.0.3), with results that did not survive correction explicitly noted as exploratory findings.

## 3. Results

### 3.1 Anticipatory pleasure moderates the relationship between reward sensitivity and behavioral motivation

Our first aim was to examine the relationships between behavioral motivation, anticipatory pleasure (TEPSa), and reward sensitivity during reward anticipation. Behavioral motivation was indexed by reaction time (RT) contrasts in the Monetary Incentive Delay (MID) task (see Methods), with HS>LS (high stakes vs. low stakes) capturing the overall pattern of RT differentiation: faster responses to high-stakes trials (large gains or losses) compared to low-stakes trials indicate higher behavioral motivation, whereas flatter RT patterns reflect lower motivation (Figure 2A). This pattern was evident at the group level (Fig. 2B). A repeated-measures ANOVA revealed a significant effect of condition on RTs, *F*(4,180)=5.022, *p*<0.001. Post-hoc pairwise comparisons using Tukey’s HSD confirmed that participants responded faster to incentive-salient trials: RTs in the Large Gain condition were significantly lower than those in the Neutral (*t*=−3.525, *p*=0.0048) and Small Loss conditions (*t*=−2.768, *p*=0.0484), whereas Neutral RTs were significantly higher than those in the Large Loss condition (*t*=3.454, *p*=0.0061). No other pairwise comparisons reached statistical significance. Full pairwise comparisons across trial conditions are available in Supplementary Table 1. These findings demonstrate that, on average, participants exhibited greater behavioral motivation for large gain and loss trials compared to neutral trials, consistent with the motivational salience of high-stakes incentives.

We next explored how behavioral motivation (HS>LS) relates to individual differences in reward sensitivity (RS) and anticipatory pleasure (TEPSa; Fig. 2C). Contrary to our hypothesis, HS>LS was not significantly correlated with either anticipatory pleasure (*r*=0.12, *p*=0.43) or reward sensitivity (*r*=0.052, *p*=0.73). However, a significant negative correlation emerged between LG>N and RS, indicating that individuals with greater RS exhibited stronger behavioral motivation (more negative LG>N values, reflecting faster RTs) for large gain trials compared to neutral trials. Moreover, we identified a significant interaction between RS and anticipatory pleasure in predicting HS>LS (*t*(46)=2.799, *p*=0.00778), controlling for consummatory pleasure. The simple slope of RS on HS>LS was positive and significant at high TEPSa (higher anticipatory pleasure), indicating that greater RS is associated with weaker behavioral motivation, non-significant at moderate TEPSa, and negative and significant at lower TEPSa (lower anticipatory pleasure), indicating that greater RS is associated with stronger behavioral motivation (Fig. 2D).

To assess the specificity of this moderation effect to anticipatory pleasure, we examined whether consummatory pleasure similarly moderated the RS-behavior relationship. While consummatory pleasure showed a main effect on behavioral motivation (*t*=2.100, *p*=.042), it did not moderate the RS-behavior relationship. Johnson-Neyman analysis confirmed this pattern: simple slopes of RS on behavioral motivation varied significantly across levels of anticipatory pleasure but not consummatory pleasure. This dissociation supports the relevance of anticipatory-specific pleasure processes for understanding how reward sensitivity drives motivated behavior during reward anticipation.

### 3.2 Reward sensitivity and ventral striatum activation during reward anticipation (exploratory)

To assess striatal responses during reward anticipation, we extracted signal from the VS across all trial types (Fig. 3). A repeated-measures ANOVA revealed a significant main effect of condition on the measured response, *F*(4, 180)=30.84, *p*<0.001. Post-hoc pairwise comparisons using Tukey’s HSD indicated that striatal responses in the Large Gain condition were significantly higher than those in the Neutral condition (*t=10.348, p*<0.0001), the Small Loss condition (*t*=3.288, *p*=0.0105), and the Large Loss condition (*t*=2.845, *p*=0.0393). Similarly, responses in the Small Gain condition were significantly higher than those in the Neutral condition (*t*=8.351, *p*<0.0001). The Neutral condition elicited significantly lower responses compared to both the Small Loss condition (*t*=-7.060, *p*<0.0001) and the Large Loss condition (*t*=-7.503, *p*<0.0001). No significant differences were observed between the Small Gain, Small Loss, and Large Loss conditions. Full pairwise comparisons across trial conditions are available in Table 2.

**Figure 3.**
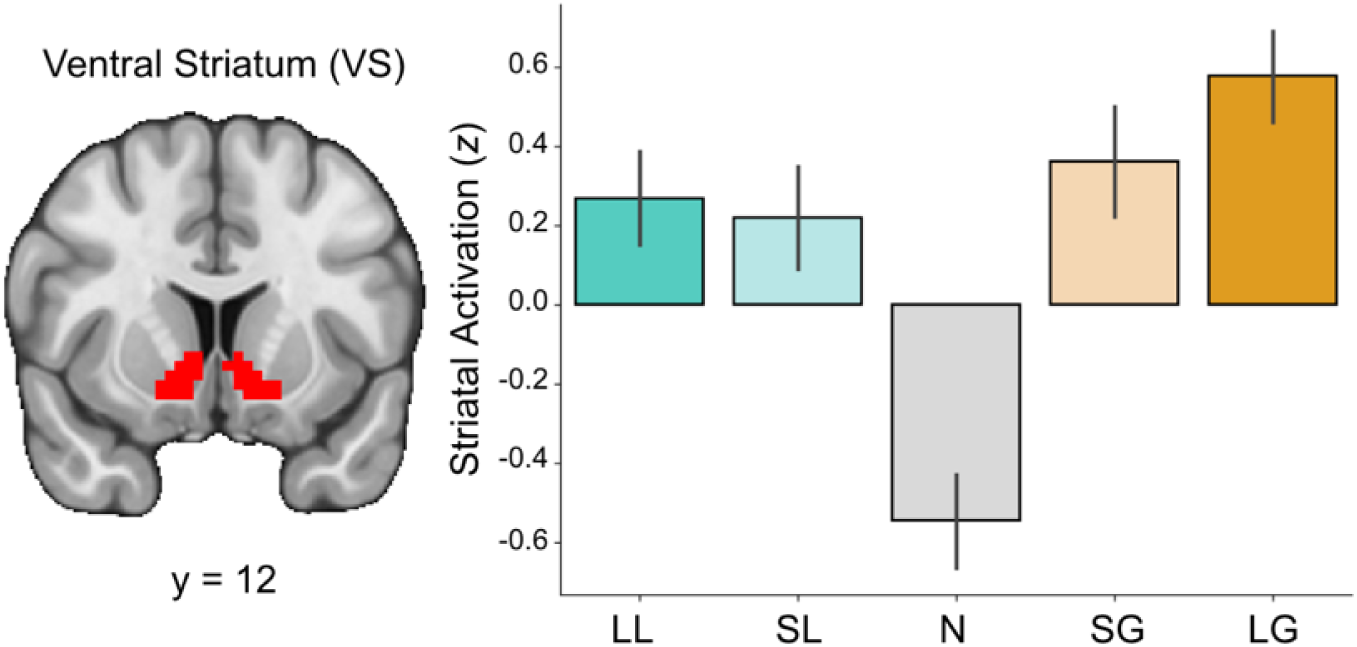
Striatal response is sensitive to incentive magnitude. Signal from the ventral striatum (VS) during the anticipation phase shows heightened activation during incentive-salient trials relative to neutral trials.

We next examined whether reward sensitivity (RS) and anticipatory pleasure (TEPSa) were associated with striatal response to varying trial types using linear regressions for the difference in striatal BOLD for task-based contrasts targeting incentive salience (i.e., HS>LS, LG>N, LL>N) with composite RS, anticipatory pleasure (TEPSa), and their interaction (RS x TEPSa) as predictors. Exploratory analyses included additional covariates for behavioral motivation. We found no significant interaction between RS and anticipatory pleasure in relation to striatal response for HS>LS (*t*=0.517, *p*=0.608), LG>N (*t*=0.923, *p*=0.361), or LL>N (*t*=-0.869, *p*=0.390). Exploratory analyses of lower order effects also did reveal that aberrant RS (i.e., reward hypo- and hyper-sensitive individuals) was associated with striatal response to large vs. small gains, such that aberrant RS was associated with reduced striatal activation relative to moderate reward sensitivity (*t*=2.368, *p*=0.023), although these did not survive correction for multiple comparisons. Full comparisons are available in Supplementary Table 2.

### 3.3 Behavioral motivation moderates the relationship between reward sensitivity and DMN-VS connectivity (primary finding)

Our final analyses examined DMN-VS connectivity using ROI-based network psychophysiological interaction (nPPI) analysis. We conducted linear regressions for the difference in connectivity for task-based contrasts targeting incentive salience (i.e., HS>LS, LG>N, Large LL>N), regressing these contrasts onto models of composite RS, Behavioral Motivation (matching the RT contrast to the PPI contrast), and their interaction (RS x Behavioral Motivation). We also tested models of RS x TEPSa and Behavioral Motivation x TEPSa.

Significant interactions between RS and Behavioral Motivation were observed across all three reward contexts (HS>LS: *β*=-12.051154, *SE*=4.689934, *t*=-2.570, *p*=0.0138; LG>N: *β*=-1.72548, *SE*=0.61087, *t*=-2.825, *p*=0.00721; LL>N: *β*=1.316432, *SE=*0.475776, *t*=2.767, *p*=0.00838; Fig. 4). For HS>LS, the simple slope of RS on DMN-VS connectivity was positive and significant at greater (i.e., more negative) behavioral motivation, indicating that among individuals with high sensitivity to HS>LS, greater RS was associated with enhanced DMN-VS connectivity. The slope was non-significant at moderate behavioral motivation, suggesting no consistent relationship at average motivation levels. The slope was negative and significant at lower (i.e., more positive) behavioral motivation, indicating that among individuals with lesser sensitivity to HS>LS, greater RS was associated with blunted DMN-VS connectivity. For LG>N (Supplementary Fig. 1), a similar pattern was observed, with positive and significant slopes at greater behavioral motivation, non-significant slopes at moderate levels, and negative slopes at lesser behavioral motivation. For LL>N (Supplementary Fig. 2), this pattern diverges: the slope is negative and significant at high levels of behavioral motivation and non-significant at lower levels, indicating blunted connectivity for individuals with greater sensitivity to LL>N and no relationship at moderate or higher levels. These effects are all preserved when controlling for TEPSa (HS>LS: *p*=0.0150; LG>N: *p*=0.00798; LL>N: *p*=0.00867). No significant interaction effects with TEPSa were observed.

**Figure 4.**
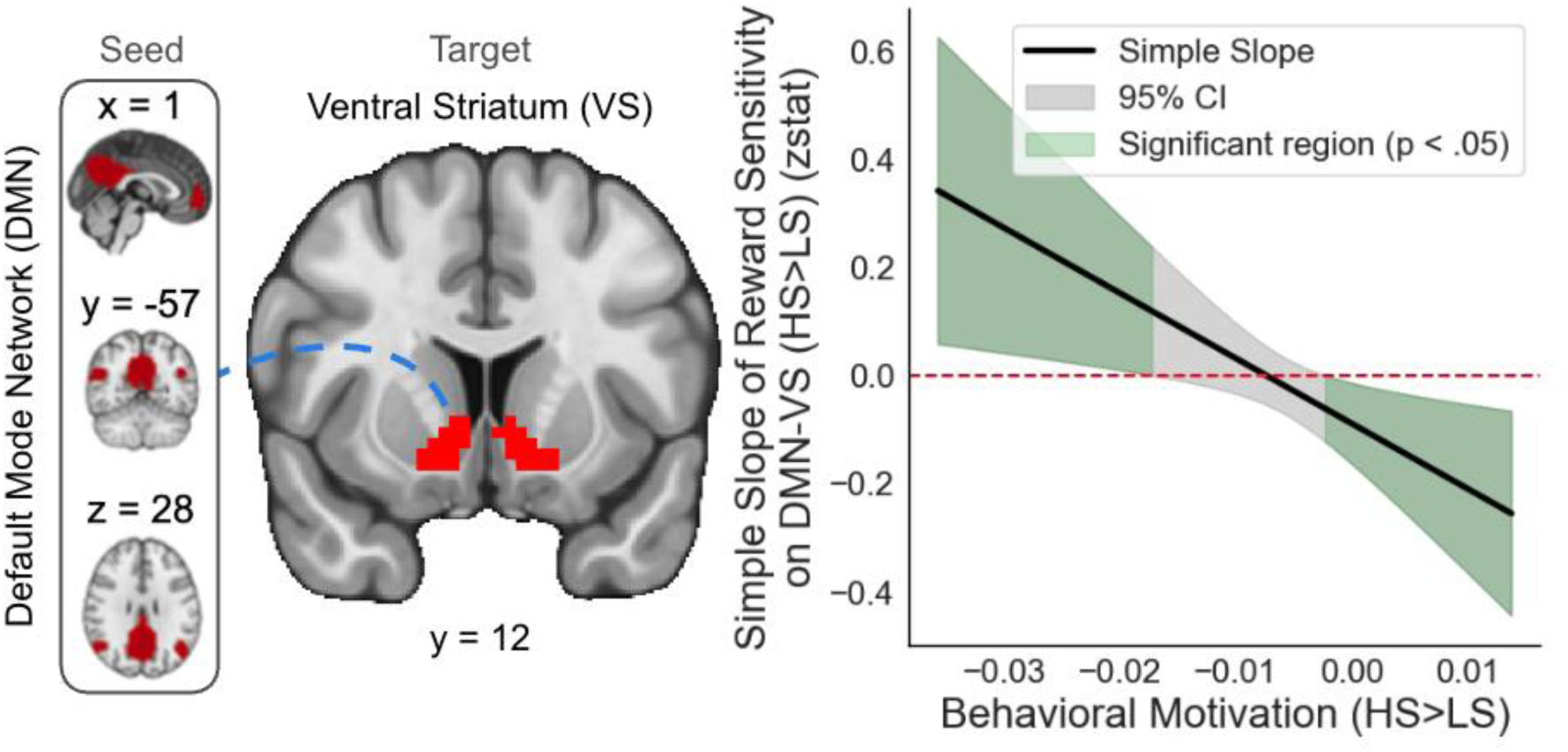
Behavioral motivation modulates the relationship between reward sensitivity (RS) and DMN-VS connectivity during reward anticipation. Johnson-Neyman plot showing the simple slope of RS and default mode network-ventral striatum (DMN-VS) connectivity for high-stakes vs. low-stakes (HS>LS) trials across behavioral motivation. Green regions indicate significant slopes (p<.05) at lower and higher behavioral motivation; the gray region indicates non-significant slopes. Among individuals with higher or lower behavioral motivation, greater RS is associated with enhanced or blunted DMN-VS connectivity, respectively.

## 4. Discussion

This study provides new evidence that behavioral motivation moderates the relationship between trait reward sensitivity and default mode network-ventral striatum (DMN-VS) connectivity during reward anticipation, with distinct patterns across high-stakes, large gain, and large loss contexts. Our study revealed complex interactions between trait reward sensitivity, behavioral motivation, and anticipatory pleasure that were correlated with neural and behavioral responses during reward anticipation. Behaviorally, participants demonstrated significant modulation of reaction times based on reward salience, responding faster for high-stakes trials relative to low-stakes trials. At the neural level, ventral striatum activation was significantly modulated by incentive magnitude, with highest activation observed for large gain trials. These findings persisted after controlling for anticipatory pleasure, suggesting a robust neurobiological mechanism underlying individual differences in reward processing.

These findings align with existing literature on the neural basis of reward processing (Knutson et al., 2001; Haber & Knutson, 2010) and extend our network integration framework of reward processing. Our observation that ventral striatal activation is modulated by incentive magnitude during the anticipation phase replicates previous MID research (Oldham et al., 2018; Büchel et al., 2017), whereas our behavioral findings showing faster reaction times for high-stakes trials support motivational intensity theory (Berridge & Robinson, 2016). The interaction between reward sensitivity and anticipatory pleasure in predicting behavioral motivation demonstrates that the translation of trait reward sensitivity into motivated behavior depends critically on anticipatory pleasure capacity. Specifically, individuals with higher anticipatory pleasure showed weaker behavioral motivation when reward sensitivity was greater, while those with lower anticipatory pleasure showed stonger behavioral motivation when reward sensitivity was greater, suggesting that anticipatory pleasure capacity fundamentally alters trait-to-behavior coupling. The specificity of this moderation to anticipatory pleasure (rather than consummatory pleasure) aligns with the MID task’s focus on anticipatory processes. When both TEPS subscales were examined, only anticipatory pleasure moderated the RS-behavior relationship, while consummatory pleasure showed a general main effect without modulation. This dissociation supports theoretical models distinguishing anticipatory (’wanting’) from consummatory (’liking’) reward components (Berridge & Robinson, 2003) and demonstrates that anticipatory pleasure capacity specifically shapes how trait reward sensitivity manifests in motivated behavior during reward anticipation.

The present study makes several novel contributions by demonstrating that behavioral motivation systematically moderates the relationship between reward sensitivity and DMN-VS connectivity during reward anticipation. Rather than a uniform relationship, we find that for individuals with higher behavioral motivation, reward sensitivity is associated with stronger DMN-VS connectivity, whereas for those with lower motivation, reward sensitivity is associated with weaker connectivity. This pattern suggests that motivational context is associated with differences in how trait sensitivity manifests through neural network integration. When motivation is high, reward-sensitive individuals show stronger coupling between self-referential processing (DMN) and reward evaluation (VS) systems, potentially reflecting enhanced integration of personal goals with incentive information. Conversely, when motivation is low, even highly reward-sensitive individuals show diminished DMN-VS integration, highlighting how state-level factors can modulate trait-level tendencies in neural processing (Insel et al., 2017; Harsay et al., 2011; Murty et al., 2017). Our findings highlight the importance of considering both trait and state factors when examining reward anticipation, as reward sensitivity effects on corticostriatal connectivity are contingent upon motivational state. This perspective may help reconcile discrepancies in previous research by identifying behavioral motivation as a key moderating factor that previous studies may have overlooked.

These findings support our network integration model of reward processing, where individual differences in reward sensitivity manifest through coordinated activity between brain systems rather than isolated regional responses. The differential patterns across gain and loss contexts suggest distinct neural mechanisms for approach versus avoidance motivation, consistent with prospect theory’s emphasis on loss aversion (Kahneman & Tversky, 1979). This framework may help reconcile previous inconsistencies in reward processing literature by identifying behavioral motivation as a key moderating factor that determines when enhanced or reduced connectivity emerges. Future research should investigate whether context-dependent modulation of DMN-VS connectivity across gain- and loss-based motivation serves as a neurobiological mechanism underlying motivational impairments in disorders such as anhedonia, depression, or apathy-related syndromes (Treadway, 2015; Roiser & Husain, 2023).

These results have implications for understanding motivational disorders. In depression, where deficits in anticipatory pleasure and motivation are central (Treadway & Zald, 2011), altered DMN-VS connectivity patterns may represent a key neural mechanism underlying anhedonia. The failure to appropriately modulate network connectivity based on motivational context could impair reward-seeking behavior and contribute to amotivation. Similarly, in addiction, disrupted DMN-VS connectivity may contribute to impaired reward-based decision-making (Volkow et al., 2010). Understanding how motivational context shapes trait-network relationships could inform personalized treatments targeting specific neural and motivational profiles.

Several limitations of the current study warrant consideration. First, our sample size (N=48) was limited due to constraints imposed by the COVID-19 pandemic, which necessitated early termination of data collection. Replication in a larger, more diverse sample would strengthen the generalizability of our findings. Second, our predominantly female sample (77.3%) also may affect generalizability, particularly given documented sex differences in reward processing (Dreher et al., 2007). Future studies should include sex as a covariate and ensure more balanced representation. Third, because behavioral motivation and neural responses were measured during the same task epoch, our design cannot separate motivational processes from other overlapping components of the MID (e.g., motor preparation and execution). As a result, connectivity differences could partially reflect behavioral performance rather than purely underlying motivational mechanisms. Our imaging models did not include RT as a covariate, and although behavioral motivation was not correlated with average RT, we cannot rule out shared variance. Future studies should incorporate independent measures of motivation (e.g., effort-based paradigms such as Treadway et al.’s effort expenditure for rewards task) or include behavioral covariates to better distinguish neural from performance-related contributions. Moreover, future studies would benefit from multi-method approaches combining self-report, behavioral, and physiological measures to more comprehensively assess hedonic capacity and better distinguish trait-like from state-like components of anhedonia. Our non-clinical sample likely experienced less mood-related fluctuation than clinical samples, but replication in clinical samples would strengthen construct validity. Fourth, although the MID task effectively captures anticipatory processes, it represents only one facet of reward processing; future studies should employ multiple paradigms to dissociate anticipation, consumption, and learning components, which engage overlapping but distinct neural circuits (Wang et al., 2016; Haber & Knutson, 2010). Additionally, the differential connectivity patterns for gains versus loss anticipation suggest separate neural mechanisms for approach versus avoidance motivation that merit further investigation (Supplementary Figs. 1 and 2; Aupperle et al., 2015; Spielberg et al., 2012). Finally, examining how these findings relate to problematic substance use remains an important avenue for future research, as altered reward sensitivity and motivation have been implicated in addiction vulnerability (Volkow et al., 2010).

In conclusion, our study highlights the value of integrating behavioral, self-report, and neuroimaging measures to develop a more comprehensive model of reward processing. This approach may help us better understand the complex interplay between stable traits (such as reward sensitivity) and context-dependent states (such as current levels of motivation) in clinical populations (Pizzagalli, 2014). Ultimately, these insights may inform more personalized treatments for conditions like depression and addiction, where reward dysfunction is a central feature, by tailoring interventions based on an individual’s specific neural and motivational profiles.

## 5. Declarations

### Ethics approval

This research was conducted in compliance with the Declaration of Helsinki code of ethics and the standards established by the Temple University Institutional Review Board.

### Consent to participate

Written informed consent was obtained from all participants (all aged 18+).

### Consent for publication

Not applicable.

### Availability of data and materials/Open practices statement

The data and materials for this experiment are available at OpenNeuro Dataset 4920 (Smith et al., 2024).

### Code availability

Analytic code will be made available on GitHub: https://github.com/DVS-Lab/istart-mid-clean. This study was pre-registered: https://aspredicted.org/PQA_WPB.

### Authors’ contributions

All authors meet the criteria for authorship and approved the manuscript for submission.

