## Supplementary Information for "Trait reward sensitivity and behavioral motivation are associated with connectivity between the default mode network and the striatum during reward anticipation" (from another version of this preprint: version 3, 2025-09-10)

**Supplementary Methods**

**SM1. *Deviations from Pre-Registration***

Whereas our pre-registration outlined an analysis of reaction time (RT) patterns across trial types to assess behavioral motivation, we later expanded our approach to include specific contrasts for large gain > neutral (LG > N) and large loss > neutral (LL > N). These contrasts were added to more directly capture the impact of reward and loss outcomes on behavioral motivation, given prior evidence that both large rewards and large losses significantly influence motivation (Kahneman & Tversky, 1979). Although these contrasts were not pre-registered, they were deemed essential for a more precise analysis of reward salience and were fully aligned with the study's core aims.

**SM2. *Behavioral Analyses***

For analyses involving pairwise comparisons between trial conditions, we conducted a total of 10 tests (comparing each of the five conditions—Large Gain, Small Gain, Neutral, Small Loss, and Large Loss—against every other condition). To control for the family-wise error rate and minimize Type I errors, we applied Tukey's Honest Significant Difference (HSD) correction for all pairwise post-hoc comparisons following significant omnibus ANOVA results. For our exploratory analyses examining the relationships between individual difference measures (reward sensitivity, anhedonia) and neural responses, we report both uncorrected p-values and whether findings survived correction for multiple comparisons. Specifically, for analyses investigating VS activation across different reward contrasts, we considered a family of 3 tests (incentive salience contrasts, HS>LS, LG>N, LL>N) and applied a Bonferroni-corrected significance threshold of p < 0.0167 (0.05/3). Similarly, for our analyses of DMN-VS connectivity, we considered the same family of 3 contrasts and interaction terms with individual difference measures, applying the same correction threshold. All statistical analyses were performed using R (version 4.0.3; R Core Team, 2020), and results that did not survive multiple comparison correction are explicitly noted as exploratory findings.

**SM3. *Neuroimaging Data Acquisition and Preprocessing***

### Neuroimaging Data Acquisition

Each run included 292 functional volumes. We also collected single-band reference images with each functional run of multi-band data to improve motion correction and registration. To facilitate anatomical localization and co-registration of functional data, a high-resolution structural scan was acquired (sagittal plane) with a T1-weighted magnetization=prepared rapid acquisition gradient echo (MPRAGE) sequence (224 mm in FOV, TR = 2,400 ms, TE = 2.17 ms, voxel size of 1.0 x 1.0 x 1.0 mm^3^, flip angle 8°). In addition, we also collected a B0 fieldmap to unwarp and undistort functional images (TR: 645 ms; TE1: 4.92 ms; TE2: 7.38 ms; matrix 74×74; voxel size: 2.97×2.97×2.80 mm^3^; 58 slices, with 15% gap; flip angle: 60°)

### Pre-processing of Neuroimaging Data

#### Anatomical data pre-processing

The T1-weighted (T1w) image was corrected for intensity non-uniformity (INU) with `N4BiasFieldCorrection`, distributed with ANTs 2.3.3, and used as T1w-reference throughout the workflow. The T1w-reference was then skull-stripped with a *Nipype* implementation of the `antsBrainExtraction.sh` workflow (from ANTs), using OASIS30ANTs as target template. Brain tissue segmentation of cerebrospinal fluid (CSF), white-matter (WM), and gray-matter (GM) was performed on the brain-extracted T1w using `fast` (FSL 5.0.9). Volume-based spatial normalization to one standard space (MNI152NLin2009cAsym) was performed through nonlinear registration with `antsRegistration` (ANTs 2.3.3), using brain-extracted versions of both T1w reference and the T1w template. The following template was selected for spatial normalization: *ICBM 152 Nonlinear Asymmetrical template version 2009c* (TemplateFlow ID: MNI152NLin2009cAsym)

#### Functional data pre-processing

Primarily, for each of the BOLD runs per subject, the following pre-processing steps were performed. First, a reference volume and its skull-stripped version were generated by aligning and averaging 1 single-band references (SBRefs). A B0- nonuniformity map (or *field map*) was estimated based on a phase-difference map calculated with a dual-echo GRE (gradient-recall echo) sequence, processed with a custom workflow of *SDCFlows* inspired by the `epidewarp.fsl` script, (<http://www.nmr.mgh.harvard.edu/~greve/fbirn/b0/epidewarp.fsl>) and further improvements in HCP Pipelines. The *field map* then was co-registered to the target EPI (echo-planar imaging) reference run and converted to a displacements field map (amenable to registration tools such as ANTs) with FSL's `fugue` and other *SDCflows* tools. Based on the estimated susceptibility distortion, a corrected EPI (echo-planar imaging) reference was calculated for a more accurate co-registration with the anatomical reference. The BOLD reference was then co-registered to the T1w reference using `flirt` (FSL 5.0.9) with the boundary-based registration cost-function. Co-registration was configured with nine degrees of freedom to account for distortions remaining in the BOLD reference. Head-motion parameters with respect to the BOLD reference (transformation matrices, and six corresponding rotation and translation parameters) are estimated before any spatiotemporal filtering using `mcflirt`. Further, we applied spatial smoothing with a 5mm full-width at half-maximum (FWHM) Gaussian kernel using FEAT (FMRI Expert Analysis Tool) Version 6.00, part of FSL (FMRIB’s Software Library, www.fmrib.ox.ac.uk/fsl). Non-brain removal using BET (Smith, 2002) and grand mean intensity normalization of the entire 4D dataset by a single multiplicative factor were also applied.

BOLD runs were slice-time corrected using `3dTshift` from AFNI 20160207. First, a reference volume and its skull-stripped version were generated using a custom methodology of *fMRIPrep*. The BOLD time-series (including slice-timing correction when applied) were resampled onto their original, native space by applying a single, composite transform to correct for head-motion and susceptibility distortions. These resampled BOLD time-series will be referred to as *preprocessed BOLD in original space*, or just *preprocessed BOLD*. The BOLD time-series were resampled into standard space, generating a *preprocessed BOLD run in MNI152NLin2009cAsym space*. First, a reference volume and its skull-stripped version were generated using a custom methodology of *fMRIPrep*. Several confounding time-series were calculated based on the *preprocessed BOLD,* notably including framewise displacement (FD).

Additionally, a set of physiological regressors were extracted to allow for component-based noise correction (*CompCor*). These components are estimated after high-pass filtering; the *preprocessed BOLD* time-series (using a discrete cosine filter with 128s cut-off) for anatomical component correction (aCompCor). For aCompCor, three probabilistic masks (CSF, WM and combined CSF+WM) are generated in anatomical space. The implementation differs from that of Behzadi et al. in that instead of eroding the masks by 2 pixels on BOLD space, the aCompCor masks are subtracted from a mask of pixels that likely contain a volume fraction of GM. This mask is obtained by thresholding the corresponding partial volume map at 0.05, and it ensures components are not extracted from voxels containing a minimal fraction of GM. Finally, these masks are resampled into BOLD space and binarized by thresholding at 0.99 (as in the original implementation). Components are also calculated separately within the WM and CSF masks. For each CompCor decomposition, the *k* components with the largest singular values are retained, such that the retained components' time series are sufficient to explain 50 percent of variance across the nuisance mask (CSF, WM, combined, or temporal). The remaining components are dropped from consideration. The head-motion estimates calculated in the correction step also were placed within the corresponding confounds file. All resamplings can be performed with a single interpolation step by composing all the pertinent transformations (i.e., head-motion transform matrices, susceptibility distortion correction when available, and co-registrations to anatomical and output spaces). Gridded (volumetric) resamplings were performed using `antsApplyTransforms` (ANTs), configured with Lanczos interpolation to minimize the smoothing effects of other kernels.

Many internal operations of *fMRIPrep* use *Nilearn* 0.6.2, mostly within the functional processing workflow. For more details of the pipeline, see the section corresponding

to workflows in *fMRIPrep*'s documentation (https://fmriprep.readthedocs.io/en/latest/workflows.html).

**Supplementary Table 1. *Pairwise comparisons of reaction times across trial conditions.***

| Contrast | Estimate | *t* | *p* |
| --- | --- | --- | --- |
| Large Gain - Small Gain | -0.007629 | -1.278 | 0.705 |
| Large Gain - Neutral | -0.02104 | -3.525 | 0.0048** |
| Large Gain - Small Loss | -0.016521 | -2.768 | 0.0484* |
| Large Gain - Large Loss | -0.000424 | -0.071 | 1 |
| Small Gain - Neutral | -0.013411 | -2.247 | 0.1675 |
| Small Gain - Small Loss | -0.008892 | -1.49 | 0.5705 |
| Small Gain - Large Loss | 0.007205 | 1.207 | 0.7473 |
| Neutral - Small Loss | 0.004519 | 0.757 | 0.9424 |
| Neutral - Large Loss | 0.020616 | 3.454 | 0.0061** |
| Small Loss - Large Loss | 0.016098 | 2.697 | 0.0584 |
| *(p<.05); **(p<.01). | |  |  |

**Supplementary Table 2. *Pairwise comparisons of striatal response to incentive magnitude.***

| Contrast | Estimate | *t* | *p* |
| --- | --- | --- | --- |
| Large Gain - Small Gain | 0.216 | 1.996 | 0.272 |
| Large Gain - Neutral | 1.121 | 10.348 | <.0001*** |
| Large Gain - Small Loss | 0.356 | 3.288 | 0.0105* |
| Large Gain - Large Loss | 0.308 | 2.845 | 0.0393* |
| Small Gain - Neutral | 0.905 | 8.351 | <.0001*** |
| Small Gain - Small Loss | 0.14 | 1.292 | 0.6966 |
| Small Gain - Large Loss | 0.092 | 0.849 | 0.9149 |
| Neutral - Small Loss | -0.765 | -7.06 | <.0001*** |
| Neutral - Large Loss | -0.813 | -7.503 | <.0001*** |
| Small Loss - Large Loss | -0.048 | -0.443 | 0.992 |
| *(p<.05); ***(p<.001). | |  |  |

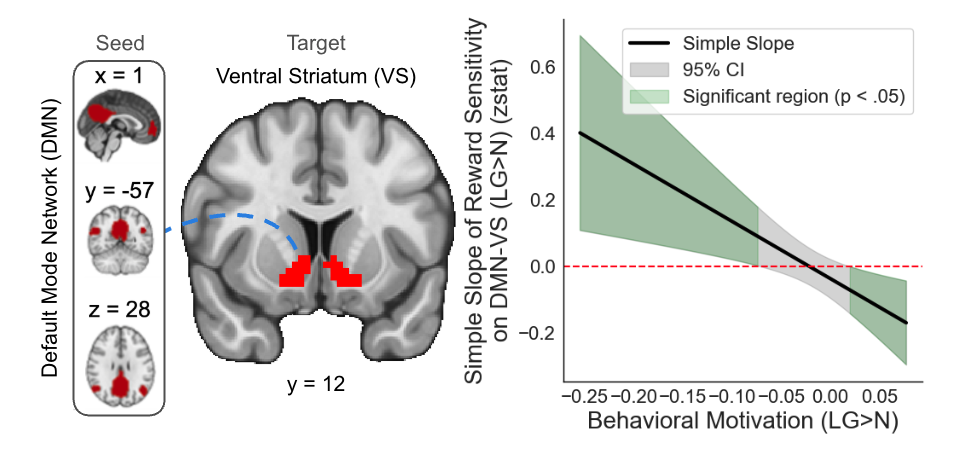

**Supplementary Figure 1. *Behavioral motivation (LG>N) modulates the relationship between reward sensitivity (RS) and DMN-VS connectivity during reward anticipation.*** *Johnson-Neyman plot showing the simple slope of RS and default mode network-ventral striatum (DMN-VS) connectivity for large gain vs. neutral (LG>N) trials across behavioral motivation. Green regions indicate significant slopes (p<.05) at lower and higher behavioral motivation; the gray region indicates non-significant slopes. Among individuals with higher (i.e., more negative values) or lower (i.e., more positive) behavioral motivation, greater RS is associated with enhanced or blunted DMN-VS connectivity, respectively.*

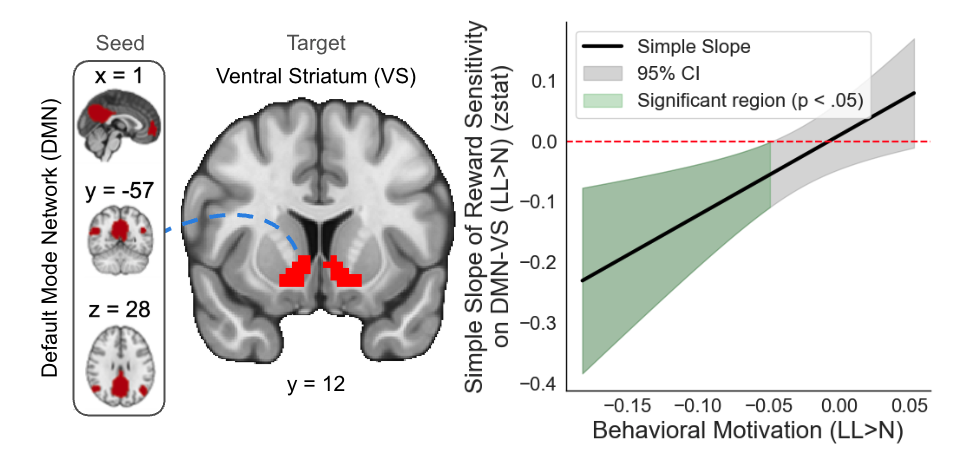

**Supplementary Figure 2. *Behavioral motivation (LL>N) modulates the relationship between reward sensitivity (RS) and DMN-VS connectivity during reward anticipation.*** *Johnson-Neyman plot showing the simple slope of RS and default mode network-ventral striatum (DMN-VS) connectivity for large loss vs. neutral (LL>N) trials across behavioral motivation. The green region indicates a significant negative slope (p<.05) at lower behavioral motivation; the gray region indicates non-significant slopes. Among individuals with higher behavioral motivation for large losses (i.e., more negative values), greater RS is associated with blunted DMN-VS connectivity.*
